# *DELAY OF GERMINATION 6*, encoding the *ANAC060* transcription factor, inhibits seed dormancy

**DOI:** 10.1101/2021.05.03.442418

**Authors:** Shuang Song, Hanzi He, Kerstin Gühl, Marieke van Bolderen-Veldkamp, Gonda Buijs, Leo A.J. Willems, Leónie Bentsink

**Affiliations:** Laboratory of Plant Physiology, Wageningen University, 6708 PB Wageningen, The Netherlands; College of Plant Science and Technology, Huazhong Agricultural University, Wuhan 430070, Hubei, China; Department of Plant Biology, University of Geneva, 1211 Geneva Switzerland

**Keywords:** Seed dormancy, *ANAC060*, PP2CAs, *DELAY OF GERMINATION 6 (DOG6)*, Membrane binding domain

## Abstract

The timing of seed germination is regulated by seed dormancy. There is ample natural variation for seed dormancy among as well as within plant species. In Arabidopsis several *DELAY OF GERMINATION* quantitative trait loci have been identified, of which *DOG1* is best studied. Here we report the identification of *DOG6*, a quantitative trait locus with a similar strong effect on seed dormancy as *DOG1. DOG6* affects the timing of germination both in laboratory as well as in field conditions. Complementation cloning revealed that *DOG6* encodes the membrane bound transcription factor *ANAC060*. The absence of the ANAC060 protein or its sequestration outside the nucleus results in increased seed dormancy levels. The different natural variants of *ANAC060* differ for the presence of the membrane binding domain, either due to the fact that this domain is absent in the genomic sequence or because the cDNA is alternatively spliced. Our data indicates that ANAC060 regulates seed dormancy by among others binding to and regulating the expression of protein phosphatases 2C class A proteins including PROTEIN PHOSPHATASE 2CA (PP2CA), ABI FIVE BINDING PROTEIN 3 (AFP3) and HIGHLY ABA-INDUCED PP2C GENE 3 (HAI3).

**Significance Statement:** ANAC transcription factors are known to effect plant development as well as the response of plants to their environment. Here, we present the identification of *DELAY OF GERMINATION 6* (*DOG6*), a seed dormancy quantitative trait locus that encodes the *ANAC060* transcription factor. We have identified different natural alleles of *ANAC060* and show that these genetic variants determine the localization of the protein. *ANAC060* alleles that lack the membrane binding domain end up in the nucleus. Hence they affect transcription and as such attenuate seed dormancy.

## Introduction

The timing of seed germination is essential for the survival of seed plants. This timing of germination is controlled by seed dormancy. Seed dormancy is an important life history trait for which ample genetic variation is present in nature. Studies employing this natural variation in *Arabidopsis thaliana* European accessions have revealed that low altitudes, high latitudes, high temperature, low summer precipitation and high radiation correlate with high seed dormancy levels (1, 2). The loci underlying the natural genetic variation in Arabidopsis have been identified using quantitative trait loci (QTL) mapping. In total eleven QTL have been revealed, of which two *DELAY OF GERMINATION 1* (*DOG1*) and *DOG18* (*RDO5*) have been cloned (3, 4). Mutant analyses have shown that *DOG1* functions down-stream of the ethylene receptor ETR1 and the ethylene response factor ERF12. The functioning of ERF12 depends on interaction with TOPLESS (TPL), together these proteins repress the expression of *DOG1* in the presence of ethylene (5). Moreover, *DOG1* expression is known to be increased by cold (6). Bryant *et al*. (7) showed that LEUCINE ZIPPER TRANSCRIPTION FACTOR67 (bZIP67) protein increases in abundance when seeds perceive cold during maturation and that this transactivates *DOG1* and thereby helps to establish primary dormancy. In mature plants *DOG1* is down regulated by its antisense transcript (*asDOG1*) (8). Inhibition by the plant hormone abscisic acid (ABA) and drought inhibits *asDOG1* expression, and therefore leads to increased *DOG1* sense transcript. The depth of dormancy correlates with the amount of DOG1 protein which loses its function during seed dry storage (after-ripening), conditions that also lead to dormancy reduction (9). Recently a model has been proposed in which DOG1 controls seed dormancy by its physical interaction with the phosphatases ABA-HYPERSENSITIVE GERMINATION 1 and 3. This interaction functionally blocks their roles in the release of seed dormancy (10). DOG18/RDO5 is a member of the protein phosphatase 2C family of that does not show phosphatase activity and is thought to act independently of DOG1 (4, 11).

Here we describe the cloning of *DOG6*, the second strongest dormancy QTL in the genetic screen in which also *DOG1* and *DOG18*/*RDO5* have been identified (12, 13). We have previously shown that several natural alleles of *DOG6* increase seed dormancy in the Landsberg *erecta* (L*er*) genetic background (13). These include the Cape Verde Islands (Cvi), Kashmir-2 (Kas-2) and Shahdara (Sha) alleles, the latter two have been used for the fine-mapping of the *DOG6* QTL. Complementation cloning revealed that *DOG6* encodes the membrane bound transcription factor *ANAC060* (*At3g44290*). We show that the natural alleles of *ANAC060* determine the dormancy behavior based on the fact of the presence of the membrane binding domain. The membrane binding domain retains the protein from the nucleus, genotypes in which ANAC060 is localized in the nucleus have a non-dormant phenotype. *ANAC060* and *DOG1* regulate seed dormancy by additive pathways, a role for *ANAC060* at the point where ABA and *DOG1* diverge has been proposed.

## Results

### *DOG6* encodes the Arabidopsis NAC transcription factor, *ANAC060*

*DOG6* has been identified as a major dormancy locus in five recombinant inbred populations that were made by crossing L*er* with other accessions. The presence of the QTL was verified using near isogenic lines (NILs) that contain introgression fragments of the Cvi, Santa Maria da Feira-0 (Fei-0), Kashmir-2 (Kas-2), Kondara (Kond) and Sha accessions at the position of the *DOG6* locus in the L*er* genetic background. All these NILs conferred a more dormant phenotype than L*er* itself (13).

To understand the molecular function of *DOG6*, the gene was cloned using positional cloning. Recombinants selected from crosses between L*er* and NIL*DOG6*-Sha and L*er* and NIL*DOG6*-Kas-2 positioned the *DOG6* locus between the markers T10D17-159697 and LK2 snp1 on chromosome 3 (Fig. 1A-D, Table S1). This region of 84.5 Kb contained 15 ORFs (Fig. 1E). Dormancy phenotyping by means of after-ripening analyses (14) on knock-out mutants of these genes located in the region identified *At3g44290* as the most likely candidate to encode for *DOG6* (Fig. S1, Table S2). Two lines, SALK_127838C and SALK_012554, with an insertion in *At3g44290* had an increased seed dormancy level when compared to wild type Col-0 (Fig. 1F, 1G). The expression of *ANAC060* was confirmed to be significantly lower in *anac060* mutants than in Col-0 (Fig. 1H). *At3g44290* encodes the membrane bound transcription factor *ANAC060*. Complementation cloning performed by cloning the *At3g44290* L*er* allele into NIL*DOG6*-Kas-2 using binary vector pKGW (pKGW-*DOG6*-L*er*) restored the reduced dormancy phenotype of L*er* (Fig. 1I), which confirmed that *DOG6* encodes the *ANAC060* transcription factor. In the following text we will refer to *ANAC060* since this is the official annotation of the gene.

**Figure 1.**
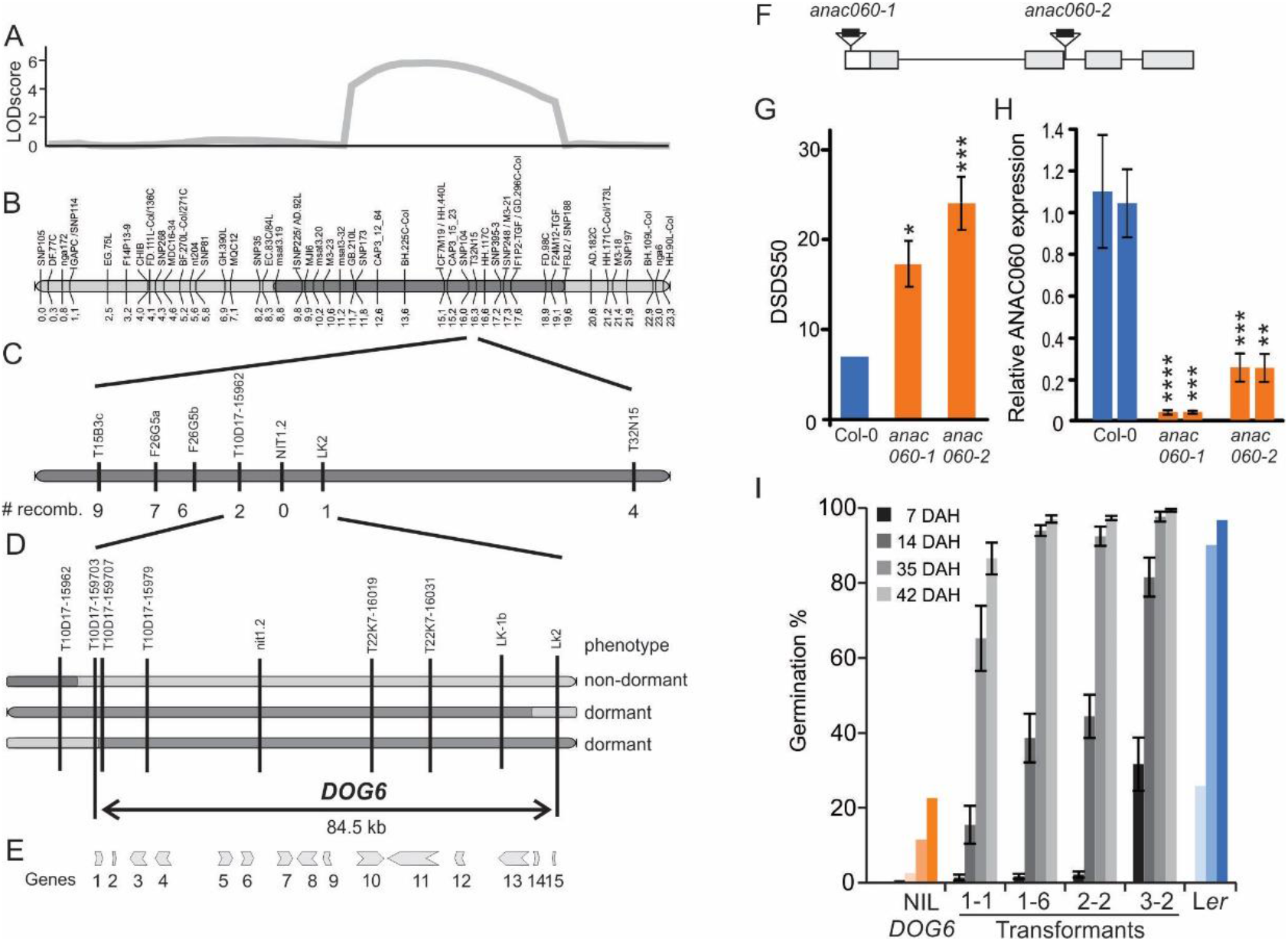
Genetic fine mapping of *DELAY OF GERMINATION 6* and the identification of the underlying gene. A. Position of the *DOG6* QTL in the genetic map of the L*er*/Kas-2 RIL population. B. Schematic representation of chromosome 3 of a near isogenic line that contains the Kas-2 introgression (dark grey bars) in the L*er* genetic background (pale grey bars). Junctions between pale and dark grey bars indicate crossover breakpoints. C. Zoom into the QTL region. D. Overview of the recombinants that are essential for determining the position of *DOG6*, at the right the dormancy phenotype of the recombinants is indicated. E. Genes that are located in the 84.5 kb fine-mapped region. As list containing the AGI numbers of the presented genes can be found in Table S2. F. schematic representation of the *ANAC060* gene model. 5’UTR in white, exons light grey and the T-DNA insertions are indicated by the black boxes. G. dormancy levels expressed DSDS50 values (Days of Seed Dry Storage required to reach 50% of germination) of the two T-DNA knock-down lines that are disrupted in the *ANAC060* gene in comparison with wild type (Col-0). Averages and SE are presented, n=9. H. Relative *ANAC060* expression in Col-0 and the two knock-down lines. The expression values were normalized using two reference genes (*At2g28390* and *At4g12590*). Averages and SE are presented, n=9. Asterisks indicate the significance determined by two-way ANOVA * P ≤ 0.05, ** P ≤ 0.01, *** P ≤ 0.001 and **** P ≤ 0.0001. I. Complementation cloning of *DOG6*/*ANAC060*. Germination behavior of NIL*DOG6*-Kas-2, four independent transformants that contain the L*er* allele of *ANAC060* in the NIL*DOG6*-Kas-2 genetic background and L*er*.

*ANAC060* belongs to OsNAC8 subgroup of group I NAC transcription factors that includes *ANAC089* and *ANAC040* (15). *ANAC060* and *ANAC089* have earlier been identified to be important for sugar sensitivity in seedlings (16, 17). We investigated the dormancy levels in knock-down mutants of these homologs and revealed that the *anac089* mutant is slightly less dormant than its wild type, whereas *anac040* mutants do not show a dormancy phenotype (Fig. S2). The group I NAC transcription factors are characterized by a N terminal NAC domain and a C-terminal alpha helix, which represents a transmembrane domain (TMD) that is known to retain the transcription factors to the cytoplasm (16, 17).

### Genetic variation for *ANAC060* determines its cellular localization and thereby seed dormancy levels

Membrane bound transcription factors are often synthesized and stored in an inactive state (bound to the membrane) and become active (release to the nucleus) only in response to the appropriate environmental signal. This allows these transcription factors to quickly respond the changed environment. Sequence analyses of the different *ANAC060* alleles (Fei-0, Kas-2, Kond, Cal, Sha, Cvi, L*er*, Col-0, An-1, Tac) revealed several polymorphisms (Fig. 2A, Table S3). The major polymorphism is a 482 bp deletion in L*er* that because of this lacks the complete fourth exon which encodes the TMD. Col-0 and An-1 embrace a SNP in the third intron, which resulted in a 20 bp extension of the fourth exon that includes an in frame TAA stop codon. This early stop codon leads to the lack of the fourth exon that contains the TMD as was earlier reported for Col-0 (17). Five other sequence nucleotide polymorphisms (SNPs) caused amino acids changes. Fei-0 contains three SNPs in the second and third exon, namely Fei-1, Fei-2 and Fei-3 respectively. Fei-1 is located in the second exon, which is at the end of NAC domain. Fei-2 and Fei-3 are in the third exon. Tac and Cvi contain SNPs in the fourth exon, which are both located in TMD. The SNP in Tac resulted in early stop codon while the SNP in Cvi caused an amino acid change.

**Figure 2.**
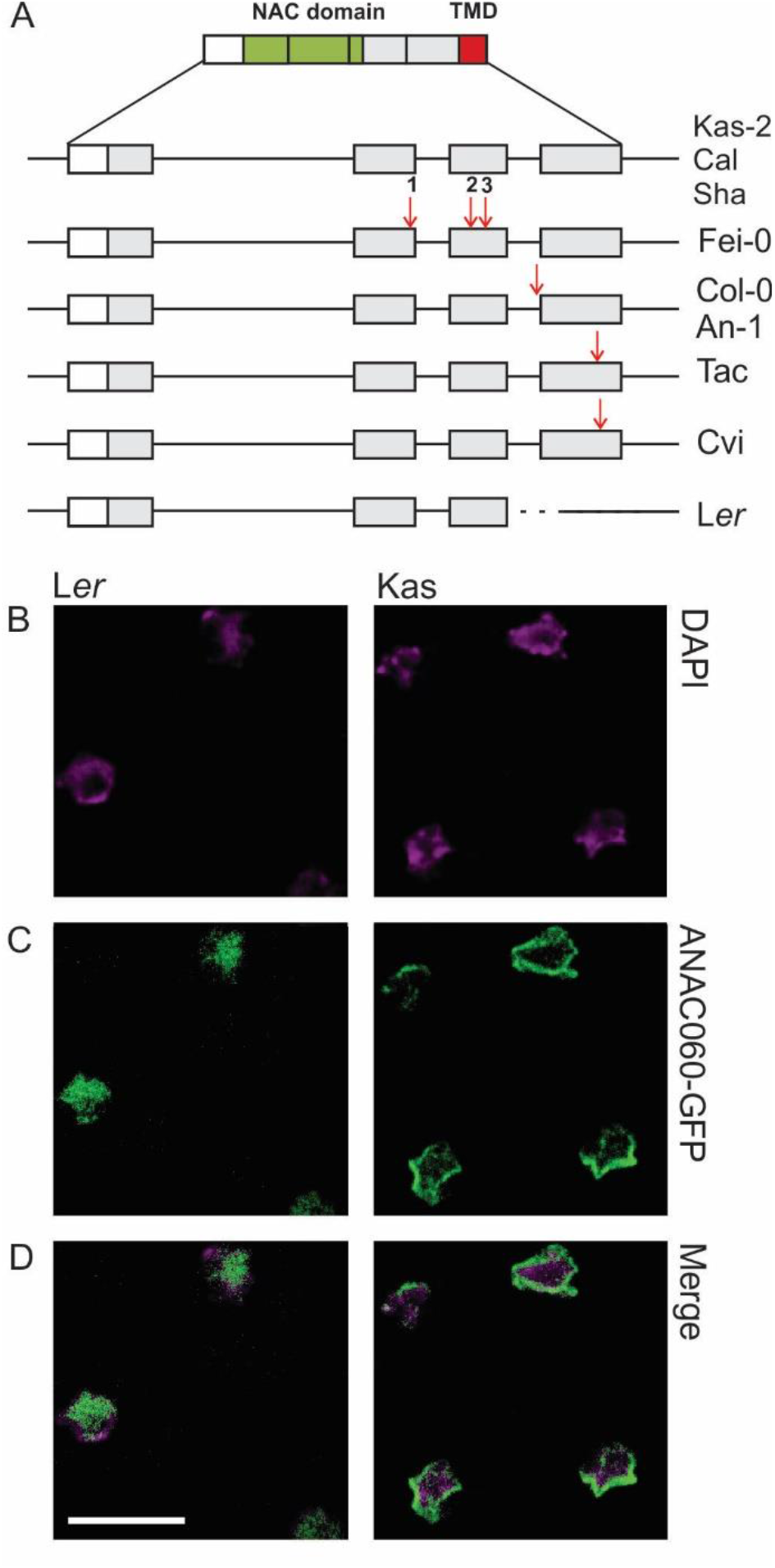
Sequence polymorphisms and cellular localization of the ANAC060 protein. A. Schematic representation of *ANAC060*. The upper panel shows the *ANAC060* cDNA with the 5’UTR (open bar), the NAC domain (green bar) and the membrane binding domain (red bar) indicated. The vertical lines indicate the exon borders. In the lower panel the polymorphisms (red arrows; nature of the polymorphisms is indicated in Table S3) of the different alleles are represented in the genomic sequence. B-D. Cellular localization of the GFP tagged *ANAC060* L*er* and *ANAC060* Kas-2 protein in the radicle of embryos of the *anac060-1* knock-down mutant, at one hour after imbibition. B. DAPI staining to indicate the location of the nucleus. C. GFP signal in seeds expressing the p*ANAC060*: GFP: *ANAC060*-L*er* and p*ANAC060*: GFP: *ANAC060*-Kas-2 fusion proteins.(D) The merge of B and C. Scale bar, 8 μm.

To investigate whether these polymorphisms affect the cellular localization of the ANAC060 protein, both transient and stable transformants were generated. *ANAC060* cDNA was fused with the green fluorescent protein (GFP) at the N-terminus with the 35S promoter for transient expression (p35S: GFP: *ANAC060*-L*er* and p35S: GFP: *ANAC060*-Kas-2) and the native Col-0 *ANAC060* promoter for stable transformed seeds (p*ANAC060*-Col-0: GFP: *ANAC060*-L*er* and p*ANAC060*-Col-0: GFP: *ANAC060*-Kas-2)(Fig. 2, Fig. S3). ANAC060-Kas-2 was located in nuclear membrane, whereas ANAC060-L*er* was solely present in the nucleus (Fig. 2B-D). Based on these and earlier findings we conclude that *ANAC060* is a transcription factor that results in a non-dormant phenotype when the protein is localized in the nucleus, a functional TMD retains the protein from the nuclear membrane which results in a higher seed dormancy. To test this hypothesis we also investigated the localization of the other alleles (i.e. the *ANAC060* Col-0, Tac, Cvi and An-1 alleles), either transiently or by stable transformants using the At2S3 seed storage protein promoter (18). These analyses confirmed the relation between a missing or mutated TMD, localization in the nucleus and a non-dormant phenotype (Fig. S3). Membrane bound transcription factors are often released from the membrane by cleavage of the TMD by membrane-associated proteases, a process referred to as intramembrane proteolysis (RIP) (19). We hypothesized that dormancy breaking cues, like after-ripening, stratification or GA treatment would trigger RIP, however we were not able to identify any GFP signal in the nucleus for any of the ANAC060 GFP tagged dormant forms (data not shown). This might be caused by the fact that RIP does not occur as well as by the instability of processed ANAC060 protein in the nucleus.

To better understand the evolutionary pressure on the SNP for *ANAC060* the rareness of these SNPs was investigated in 94 accessions that were randomly selected from a set of 349 accessions of Arabidopsis HapMap population (20). The analyses with CAPS (Cleaved Amplified Polymorphic Sequences) markers that were designed to identify the SNPs revealed that the Fei-1, Tac and Cvi mutations are rare (Table S3). The Fei-2 and Fei-3 mutations are closely linked and remarkedly three genotypes were heterozygous for the L*er* mutation.

### The Columbia allele of *ANAC060* is alternatively spliced in seeds

The expression of *ANAC060* under standard growth conditions is seed specific. The first transcripts are being detected in siliques from 8 days after pollination (DAP) onwards, expression peaks in dry mature seeds, remains high in dormant imbibed seeds and reduces in germinating seeds upon longer imbibition (Fig. 3A). As mentioned before we confirmed the presence of a SNP in the third intron of the Col-0 allele that was identified before by Li *et al*. (17). This SNP resulted in a protein lacking the TMD in seedlings grown on high sugar. However, we revealed two forms of the cDNA, one containing the extra 20 bp that results in a loss of the TMD in the protein and the other lacking the 20 bp resulting in the full length protein containing the TMD. Based on this we hypothesized that in seeds *ANAC060* is alternatively spliced (Fig. 3B). The abundance of the two cDNAs was investigated to identify whether the different forms were related to the developmental states and/or treatments that would affect the dormancy status of the seeds, however no significant differences in mRNA abundance was detected in Col-0 (Fig. 3B). The low dormancy of Col-0 is explained by the fact that the short form, which lacks the TMD, is always present. The dominance of the short form is confirmed by the dominance of the *ANAC060* L*er* allele in reciprocal crosses between NIL*DOG6* and L*er* (Fig. 3C).

**Figure 3.**
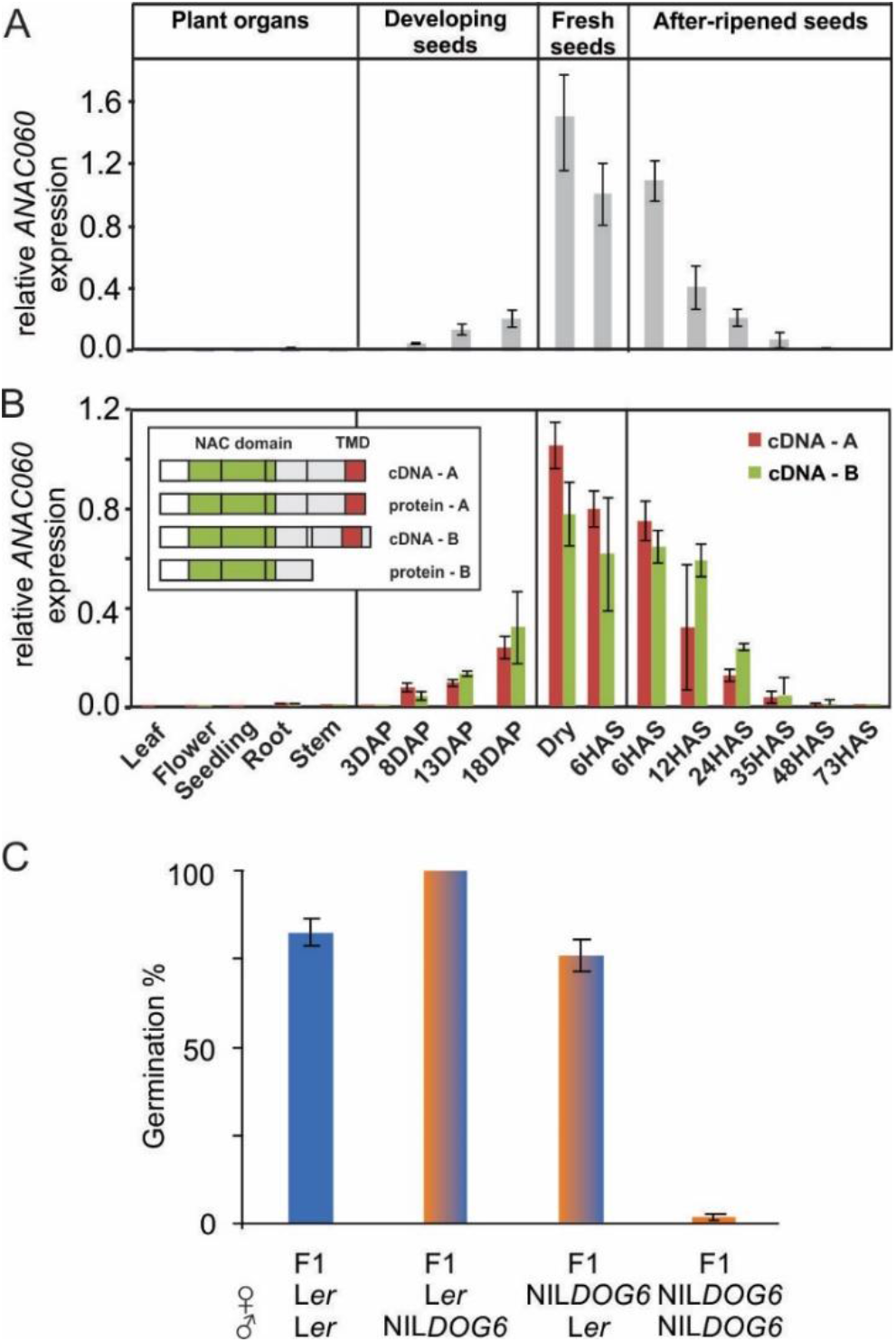
*ANAC060* alternative splicing and its germination characteristic. A. Relative *ANAC060* expression in plant organs including leaf, flower, seedling, root, stem, developing seeds, fresh seeds, after-ripened seeds. B. Two splicing forms of *ANAC060* were detected in different plant organs, leaf, flower, seedling, root, stem, developing seeds at 3, 8, 13, 18 days after pollination (DAP), freshly harvested seeds dry and 6 hours after sowing (HAS) and after-ripened dry seeds: 6, 12, 24, 35, 48, 73 HAS. Schematic presentation of the cDNAs and proteins of the two *ANAC060*-Col splicing forms is present at left top, cDNA - A refers to the annotated cDNA for which the protein contains the membrane binding domain, cDNA - B refers to the cDNA that contains 20 bp extra at the end of intron three, which results in a pre-mature stop codon and therefor a protein without the membrane binding domain. All the expression analyses have been performed on Col-0. The expression values were normalized using two reference genes (*At2g28390* and *At4g12590*) and expressed as relative values of the freshly harvested dry seeds values. n = 3 biological replicates; error bars represent SE. C. Germination behavior of L*er*, NIL*DOG6*-Cvi and their reciprocal crosses at 68 days after seed harvest.

### PROTEIN PHOSPHATASE 2C class A proteins are putative targets of *ANAC060*

*ANAC060* was described to attenuate ABA signaling in seedlings when grown on high sugar (17) and recently it was revealed that *ANAC060* directly binds to the promoter of *ABI5* and represses the sugar-induced transcription of *ABI5* (21). Chip-Seq experiments expressing *ANAC060* fused with GFP under the constitutive 35S promoter identified 500 direct targets of *ANAC060* in seedlings treated with glucose. A similar analyses in control conditions (1% sucrose) revealed 5091 putative targets (41.7% of these genes overlapped with the targets identified in the high fructose conditions). To identify targets of *ANAC060* in the regulation of seed dormancy we compared these putative targets to published gene expression data of dry and 24 imbibed dormant and fully after-ripened seeds (13, 22). We specifically addressed differences in expression between L*er* (*ANAC060* present in the nucleus and thus expected to be active as a transcription factor) and NIL*DOG6*-Kas-2 (*ANAC060* outside of the nucleus) seeds. We identified 108 genes as putative targets of *ANAC060* in the regulation of seed dormancy (Fig. 4A), of which only three genes overlap with the 500 direct targets identified in seedlings treated with glucose. This suggests that *ANAC060* regulates different genes when inhibiting seed dormancy in comparison to growth inhibition in high glucose. The three genes common between the two data sets are *At2g27300* encoding *ANAC040* (*NTM1-like 8*), *At3g23030* encoding *INDOLE-3-ACETIC ACID INDUCIBLE 2*(*IAA2*) and *At4g18650* encoding *DELAY OF GERMINATION 1-LIKE 4* (*DOGL4*). *ANAC040* is a close homologue of *ANAC060* and to investigate possible redundancy and their genetic relation we have investigated this double mutant for seed dormancy behavior. The *anac040* mutant does not show a dormancy phenotype that is different from wild type and the double mutant phenotype is indistinguishable from that of the *anac060* mutant, suggesting that *ANAC060* is epistatic over *ANAC040* (Fig. 4B). *IAA2* is an auxin responsible gene which is transiently induced by ABA in the radicle tip of the embryo before germination occurs (23). DOGL4 belongs to the DOGL family proteins but only shares a limited homology with DOG1 at the protein level. *DOGL4* is induced by ABA and plays a role in mediating reserve accumulation in seeds. *DOGL4* does not affect seed germination (24). Overall, the 106 genes are rather diverse and no specific gene ontology terms have been identified as significant (Table S4).

**Figure 4.**
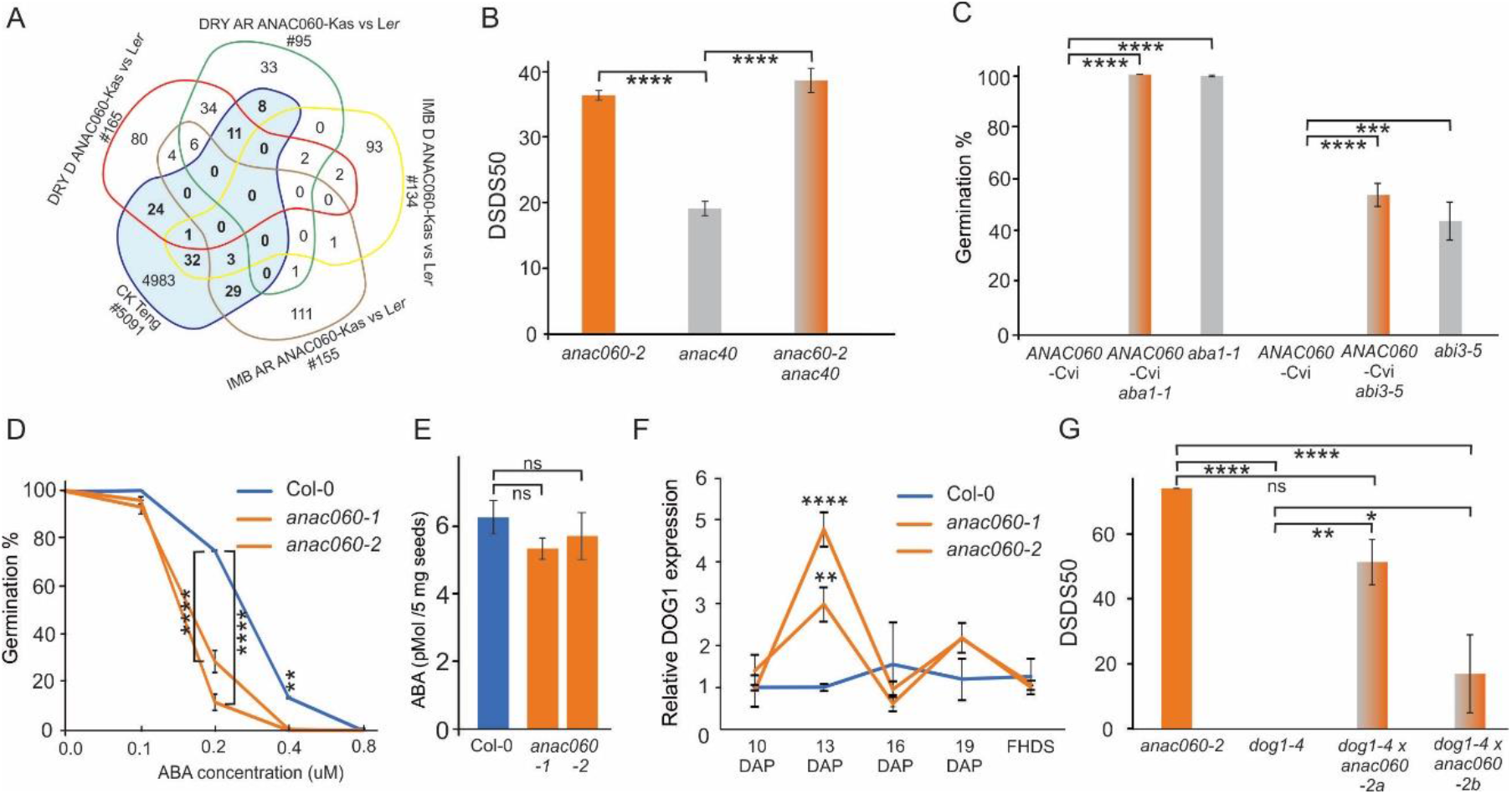
*ANAC060* putative targets. A. Venn diagram presenting the putative targets of *ANAC060*. Genes identified to directly bind to *ANAC060* in seedlings (CK Teng, (21) and those that are differentially expressed in seeds that either contain the ANAC060 protein in the nucleus or in the cytoplasm (ANAC060-L*er* compared to ANAC060-Cvi) in four different physiological stages (dry dormant seeds; DRY D, dry after ripened seeds; DRY AR, imbibed dormant seeds; IMB D and imbibed after ripened seeds; IMB AR) have been compared. In bold the 108 putative *ANAC060* targets, the gene list is provided in Table S4. B. Dormancy levels expressed as DSDS50 (Days of Seed Dry Storage required to reach 50% of germination) for *anac060-2, anac040* and the *anac060-2 anac040* double mutants. C. *ANAC060* regulated dormancy depends on ABA. Germination percentage of freshly harvested seeds of NIL*DOG6*-Cvi (*ANAC060-*Cv), *aba1-1* and the NIL*DOG6*-Cvi (*ANAC060-*Cv)*aba1-1* double mutant and NIL*DOG6*-Cvi (*ANAC060-*Cvi), *abi3-5* and the NIL*DOG6*-Cvi (*ANAC060-*Cvi)*abi3-5* double mutant. D. *ANAC060* knock-down mutants are hypersensitive to ABA. Germination behavior of Col-0 and *anac060-1* and *anac060-2* in different concentration of ABA. E. ABA levels do not differ between *ANAC060* knock-down mutants and Col-0. ABA levels in dry seeds of Col-0 and *anac060-1* and *anac060-2*. in pMol /5 mg seeds. F. Relative *DOG1* expression during seed maturation in Col-0 and *anac060-1* and *anac060-2*. RNA was extracted at 10, 13, 16 and 19 days after pollination (DAP) and of freshly harvested dry seeds (FHDS). The expression values were normalized using two reference genes (*At2g28390* and *At4g12590*). Averages and SE of four biological replicates are presented. G. *ANAC060* and *DOG1* regulate seed dormancy by independent pathways. Dormancy levels expressed as DSDS50 (Days of Seed Dry Storage required to reach 50% of germination) for *anac060-2, dog1-4* and two independent double mutants. Averages and SE of three biological replicates are presented. Asterisks indicate the significance determined by one-way ANOVA in B, C, E, G, two-way ANOVA in D, F. * P ≤ 0.05, ** P ≤ 0.01, *** P ≤ 0.001 and **** P ≤ 0.0001, ns is not significant.

**Figure 5.**
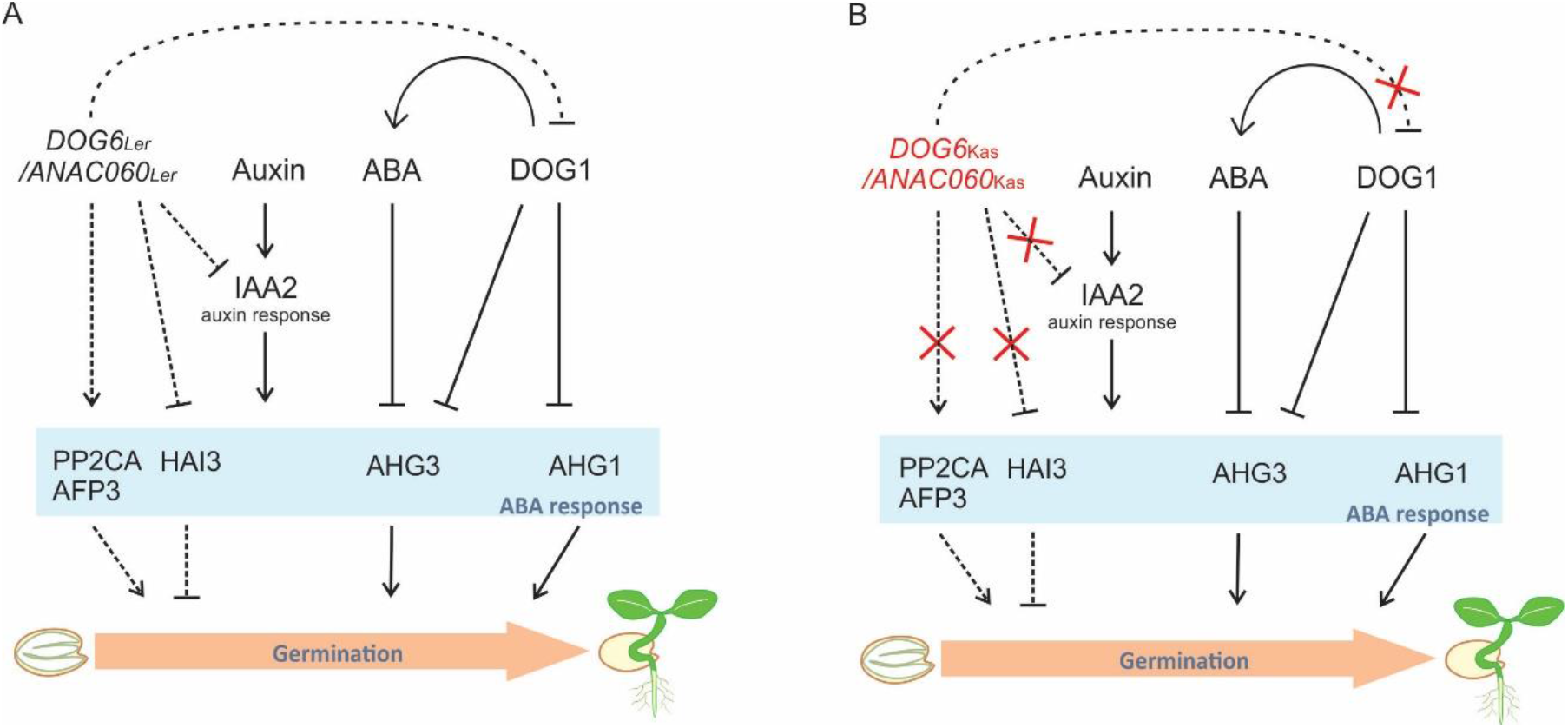
A model for the roles of *DOG6*/*ANAC060*, DOG1 and ABA in the control of seed dormancy. A. We propose that *DOG6*/*ANAC060* inhibits seed dormancy by interacting with the PP2CA class proteins (indicated in the blue box). The expression of *PP2CA* and *AFP3* is elevated in the L*er* background when compared to the *DOG6/ANAC060* Kas allele (Table S4), whereas the expression of *HAI3* is reduced in L*er*. The model is based on the active form of the ANAC060/DOG6 protein, in which the protein is present in the nucleus. PP2C proteins are essential in the regulation of seed germination. Earlier work has shown that AHG1and AHG3 are bound by DOG1 and as such play a role in the inhibition of seed dormancy by DOG1 (10). Our study revealed that DOG6/ANAC060 regulates seed germination via a pathway that is mostly independent of DOG1 (additive). The DOG6/*ANAC060* L*er* allele promotes seed germination likely by, among others, binding to the *PP2Cs*: *PP2CA, AFP3* and *HAI3*. B. This scheme presents the situation in which the *DOG6*/*ANAC060* gene is knocked-out or when the protein is associated to the membrane outside of the nucleus. The presence of ABA is absolutely required for dormancy induced by the *DOG6/ANAC060* allele that is mutated or present outside of the nucleus, however ABA levels do not differ between Col-0 and the *dog6/anac060* mutants. The *dog6/anac060* mutants are more ABA sensitive than Col-0. The effect of DOG6/ANAC060 on ABA sensitivity might either be direct of act via IAA2 that is known to affect ABA sensitivity, has been identified as a putative target of DOG6/ANAC060 and is higher expressed in dormant seeds of NILDOG6-Kas-2 when compared to L*er* (Table S4). Solid lines present proven connections based on the model presented by Nee *et al*. (10), dashed lines are based on the current study.

Not surprisingly there are also ABA related genes among the putative targets, among which *PROTEIN PHOSPHATASE 2CA* (*PP2CA*), *ABI FIVE BINDING PROTEIN 3* (*AFP3*) and *HIGHLY ABA-INDUCED PP2C GENE 3* (*HAI3*). Seeds with mutations in *PP2CA* and *AFP3* are hypersensitive to ABA, suggesting a role for *ANAC060* in overcoming ABA sensitivity which is in agreement with the earlier reported findings (17, 25-27). To further investigate the role of ABA in the regulation of seed dormancy by *ANAC060* we analyzed seed dormancy in double mutants that contained the dormant Cvi allele of *ANAC060* in the *abscisic acid insensitive 3-5* (*abi3-5*) and ABA deficient *aba1-1* genetic backgrounds. The seeds of both double mutants were completely non-dormant (Fig. 4C), confirming that the dormancy conferred by the lack of ANAC060 protein in the nucleus (*ANAC060*-Cvi allele, Fig. S3B) is ABA dependent. Moreover, the *ANAC060* knock-down mutants (*anac060-1* and *anac060-2*) are more sensitive to ABA when compared to their wild type Col-0 seeds (Fig. 4D), ABA contents do not differ between Col-0, *anac060-1* and *anac060-2* (Fig. 4E).

### *ANAC060* and *DOG1* regulate dormancy by additive pathways

*DOG1* is an important regulator of seed dormancy and although *DOG1* was not identified as a direct target of *ANAC060* we aimed at obtaining more insight into the relation between these dormancy regulators. RT-qPCR analyses during seed maturation revealed that the expression of *DOG1* is increased in the *anac060* mutant seeds at 13 days after pollination (DAP) suggesting that *ANAC060* is required for the repression of *DOG1* expression (Fig. 4F). In mature dry *anac060* seeds no differences in *DOG1* expression has been revealed when compared to Col-0 (Fig. S4). Previously, *DOG1* and *DOG6/ANAC060* have been proposed to regulate seed dormancy by additive pathways (13). To obtain more insight into the relation between these dormancy regulators the *anac060-2* mutant was crossed to the completely non-dormant *dog1-4* mutant (3). The double mutants revealed a dormancy behavior that was intermediate to that of the single mutants (Fig. 4G), which confirmed the additive effect that was reported earlier for the *DOG*NIL (NILDOG617), that contains the Cvi alleles of both *DOG1* and *DOG6*, compared to the single NILs (28).

## Discussion

The identification of genes underlying the *DOG* QTL has contributed in understanding how seed dormancy is regulated. *DOG1* was already cloned in 2006 (3), however due to its lack of conserved domains only recently its mechanisms has been being revealed (5, 7, 9, 10). With the cloning of *DOG6* the first transcription factor (*ANAC060*) underlying a *DOG* locus has been identified.

The presence of ANAC060 in the nucleus results in a non-dormant phenotype. Different genetic variants of *ANAC060* determine the localization of the protein. SNPs and deletions result in a loss of the TMD, directly or due to differential splicing. The nucleus localized ANAC060 protein, represented by its Tac and L*er* allele, is dominant. Col and An-1 alleles also result in a non-dormant phenotype due to insertion of a 22 bp fragment of the 4^th^ intron into the 4^th^ exon which results in a stop codon just before the TMD. The genetic variants of *ANAC060* can be divided into two classes structural variation already present in the genome sequence that determines whether the TMD is present or not, 2) alternative splicing, the different cDNAs in the Col and An-1 accession. The structural variants either represent point mutations that lead to early stop codons and because of that a loss of the TMD, as is the case for Tac, or a complete deletion of the TMD as is the case for the *ANAC060* L*er* allele. Also for the *ANAC060* alleles that do contain the TMD there is genetic variation that might affect the localization of the protein in the cells (Fig. S3).

*ANAC060* has previously been identified as a fructose sensing QTL in a Col/C24 F2 population (17). The Col-0 *ANAC060* allele confers sugar insensitivity and was dominant over the sugar-sensitive C24 allele. The dominant nature of the Col allele is in agreement with our findings, however we do not reveal differences in *ABSCISIC ACID INSENSITIVE 4* (*ABI4*) expression between Col and the knock-down lines (Fig. S4).

*ANAC060* belongs to a family of three transcription factors, the other members (*ANAC040* and *ANAC089*) do not significantly affect seed germination or dormancy (Fig. 4B, Fig. S2), but *ANAC040* has been identified as a putative *ANAC060* target involved in controlling both seed dormancy and sugar sensing. *ANAC040* lowers the GA threshold in response to salinity stress and thereby delays seed germination in these conditions (29). This suggests that both *ANAC040* and *ANAC060* provide adaptive mechanisms that delay seed germination in conditions that are not optimal for seedling establishment and further plant development. For *ANAC040* in vitro analyses suggest a role for paclobutrazol and cold in the processing of the protein, leading to the loss of the TMD by which the protein ends up in the nucleus (29). The effect of cold on the release of the functional NAC-TF from the plasma membrane has also been reported for *ANAC062* (30, 31). Cold stratification overcomes *ANAC060* dormancy however, we could not find any prove for post-translational regulation to be means of membrane-associated proteases that lead to loss of the TMD and nuclear localization of the ANAC060 protein (data not shown). We can however not exclude processing of the ANAC060 protein. For *ANAC089*, a close homolog of *ANAC060*, reducing conditions promote the nuclear migration (32).

NAC proteins constitute one of the largest groups of plant-specific transcription factors and are known to play essential roles in various developmental processes. They are also important in plant responses to stresses such as drought, soil salinity, cold, and heat, which adversely affect growth. Transcriptional control under environmental stresses plays a major role in plant adaptation. The importance of transcriptional regulation in plant adaptation is supported by large number of QTL which are identified based on SNPs in the promoter of the underlying gene, often resulting in mis-expression of the quantitative trait gene (33).

Quantitative trait loci analyses for seed dormancy in Arabidopsis revealed eleven *DOG* loci, for three of them the underlying genes have been identified, *DOG1* (3), *DOG18*/*RDO5* (11) and *DOG6*/*ANAC060* (current work). QTL and transcriptome analyses revealed that these QTL regulate seeds dormancy by independent pathways(13). Earlier a model has been proposed in which ABA and *DOG1* are presented as the essential regulators of seed dormancy (10). Based on the current findings we have added *ANAC060* to this model. Where the DOG1 proteins interact with ABA-HYPERSENSITIVE GERMINATION 1 (AHG1) and AHG3, *ANAC060* might interact with other PP2CAs, including PP2CA, AFP3 and HAI3. This is supported by the identification of these proteins as putative targets of *ANAC060* and the differential expression of these genes when comparing L*er* compared to NIL*DOG6*-Kas-2 (containing the ANAC060-Kas allele) seeds (Table S4).Moreover, ANAC060 affects the ABA sensitivity, this effect might either be direct or act via IAA2 which has been identified as putative target of ANAC060 and is down-regulated in L*er* compared to NIL*DOG6*-Kas-2 (containing the ANAC060-Kas allele) seeds.

## Materials and Methods

### Plant materials

*Arabidopsis thaliana* accessions Fei, Kas-2, Kond, Sha, Cvi, L*er*, Col-0, and An-1 and the T-DNA insertion lines *anac060-1*, SALK_012554(N665285); *anac060-2*, SALK_127838C (N655936); three *anac040 alleles* (N587226, N613218, N106660) and the T-DNA lines listed in Table S2 were obtained from the Arabidopsis stock centres: Arabidopsis Biological Resource Centre (ABRC) and Nottingham Arabidopsis Stock Centre (NASC). The *anac089* mutant (Gt19255) in Landsberg *erecta* (L*er*) genetic background was a gift from Dr Sheng Teng (The Chinese Academy of Sciences, Shanghai, China). The accessions Antwerp-1, Cape Verde Islands were described before (34) and Calver and Tacoma were gifts from Dr Kathleen Donohue, Duke University, NC, USA. The near isogenic lines (NILs) NIL*DOG6*-Cvi, NIL*DOG6*-Kas-2, NIL*DOG6*-Sha were described before (13). The *abscisic acid insensitive 3-5 (abi3-5)* and abscisic acid biosynthesis mutant *abscisic acid 1-1* (*aba1-1)* mutants were gifts from Dr Maarten Koornneef (Laboratory of Genetics, Wageningen University, The Netherlands). *dog1-4* (N105944; SM_3.20808) has earlier been described in (3).

### Plant growth

The plants were grown in controlled conditions 20°C/18°C (day/night) under a 16h photoperiod and 70% relative humidity using Rockwool blocks (4×4 cm) watered with Hyponex with three replicates per genotype.

### Seed germination experiments

Germination experiments were performed as described previously (35). In brief, two layers of blue germination paper were equilibrated with 48 ml demineralized water in plastic trays (15 × 21 cm). Six samples of approximately 50 to 150 seeds were spread on wetted papers using a mask to ensure accurate spacing. Piled up trays were wrapped in a closed transparent plastic bag. The experiment was carried out in a 22°C incubator under continuous light. Pictures were taken twice a day for a period of 6 d. Germination was scored using the Germinator package (34) and DSDS50 was quantified with the model described in He *et al*. (36).

### DNA and RNA isolation from seeds

#### DNA isolation

10 mg of seeds were homogenized and mixed with 520 µl extraction buffer (0.6M NaCl, 100mM Tris, 40mM EDTA, 4% Sarkosyl and 1% SDS). Heat the samples for 10 min at 65°C and put it on ice for 5 min. After centrifuging, add 600 phenol/chloroform/isoamylacohol (25:24:1) to supernatant and mix. Centrifuge again and add 0.6 volume isopropanol, then leave it for at least 10 mins at room temperature. Wash the pellet with ethanol and dry the pellet.

#### RNA isolation

Total RNA was extracted according to the hot borate protocol described by Maia, Dekkers, Provart, Ligterink and Hilhorst (37). In brief, 7-10 mg of seeds for each genotype were homogenized and mixed with 800 ml of extraction buffer containing dithiothreitol (DTT) and PVP40 which had been heated to 80°C. Proteinase K was added and incubated for 15 mins at 42°C. After adding 2M KCl, the samples were incubated on ice for 30 mins and centrifuged. Ice-cold 8M LiCl was added to the supernatant and the tubes were incubated overnight on ice. After centrifugation, the pellets were washed with ice-cold 2M LiCl and centrifuged for 10 mins. The pellets were re-suspended in 80 ml of water. The samples were phenol chloroform extracted, DNase treated and further purified. RNA quality and concentration were assessed by agarose gel electrophoresis and Nanodrop ND1000 spectrophotometry.

### Fine Mapping of *DOG6*

Details of the molecular markers used for the fine-mapping can be found in Table S1.

### Complementation of *DOG6*

Binary constructs were prepared using the Gateway Technology (Invitrogen). A genomic DNA fragment of 4.3 kb of *DOG6*-Kas was amplified by PCR using primers attB1 *DOG6* 2141-F (GGGG ACA AGT TTG TAC AAA AAA GCA GGC TTC GCTACAAGCCTACAACAATCAGACG) and attB2 *DOG6* 3’UTR-2-R (GGGG AC CAC TTT GTA CAA GAA AGC TGG GTC TCC TCC GTT AGG TTC CGT GA) and Phusion® High-Fidelity DNA Polymerase (New England Biolabs, Massachusetts, USA). This PCR product was cloned into cloned into pDONR221, resulting in pENTR-*DOG6*-Ler. pKWG *DOG6*-L*er* was produced from the above mentioned entry clone and the destination vector pKGW red seed vector (Jan Verver Molbio; Invitrogen Life Technologies) by LR reaction. pKWG *DOG6*-L*er* was introduced by electroporation into Agrobacterium tumefaciens strain GV3101, which was subsequently used to transform L*er* and NIL*DOG6*-Kas plants by floral dipping (38). Independent homozygous single insertion lines were selected based on their red fluorescence.

### Sequence analyses

*ANAC060* genetic variation was assessed by performing sequence analyses on overlapping PCR products that cover the *ANAC060* genomic region for the accessions Fei, Kas-2, Kond, Sha, Cvi, L*er*, Col-0, An-1, Cal and Tac.

### Construction of expression vectors and generation of transgenic plants

#### 35S::GFP-*ANAC060* constructs

The cDNA of Col, An, Kas-2, Cvi, Tac and L*er* was utilized to build up constructs. The cDNA of *ANAC060* was amplified with 3’ RACE-PCR (SMARTer™ RACE cDNA Amplification Kit from Clonetech) and cloned into gateway vector pDONR201. Then the cDNA was cloned into destination vector pH35NsGG which contains 35S promoter and GFP at N-terminal of *ANAC060*.

#### At2S3::GFP-*ANAC060* constructs

The cDNA of Kas-2, Cvi, Tac and L*er* was utilized to build up constructs. Firstly, the 353bp length promoter of At2S3 was cloned into gateway vector pDONR-P4P1. Then the fragment of GFP was added to the vector via pDONR207-P1P2. Then the cDNA of *ANAC060* was cloned into destination vector pDONR-P2rP3.

#### P*ANAC060* (Promoter of *ANAC060*)::GFP-*ANAC060* constructs

The native promoter of *ANAC060* (p*ANAC060*) 1746bp (17) length: Forward primer: gtatgtgagctagtctgtgagt, reverse primer: actctctttgcatcatctctaccctt was cloned into gateway vector pDONR P4P1r, the GFP fragment was added into the vector via pDONR207 P1P2, the cDNA of ANAC060-Kas and Ler were cloned in pDONRP2P3r. Three boxes were cloned into destination vector pH7m34GW R4R3.

#### DAPI staining and confocal microscopy

Seeds embryos were extracted from 1 hours imbibed seeds. The embryos were fixed with 4% PFA + 0.1% Tritonx100 in 1X PBS (pH7.4) at RT for 1 hour. This was followed by a thorough wash (4-5 times) with 1X PBS and incubated in 1ug/ml DAPI in 1X PBS for 30 mins in dark. This was followed by another wash (4-5 times) with 1X PBS. Finally, the embryos were transferred to a glass slides with 1X PBS and imaged. Confocal microscopy was conducted with Leica TCS SP8 HyD confocal microscope, with objective HC PL APO 63x/1.40 oil (DIC).

### RT-qPCR expression analyses

#### In the anac060 know down mutants

RNA was isolated from dry mature seeds as described above. cDNA was synthesized from 750ng RNA using the iScript cDNA Synthesis Kit (Bio-Rad: 170-8890) according to the manufacturer’s protocol. cDNA was diluted 10 times with sterile milliQ water. For each sample 2.5μL cDNA, 5μL iQ SYBR green super mix (Bio-Rad: 172-5125) and 0.5μL primer mix (10μL work solution) were added and supplemented with water to 10μL. RT-qPCR was performed on a CFX connect (Bio-Rad).

#### In different plant tissues

*ANAC060* expression analysis was performed using different plant tissues, including vegetative tissues (leaf (7th rosette leaf), flower (just opened flower), seedling (two weeks old seedling), root (two weeks old root), stem, developing siliques (3 days after pollination (DAP), 8DAP, 13DAP and 18DAP), dry dormant seeds and dormant seeds 6 hours after imbibition and imbibed after-ripened seeds (6h, 12h, 24h, 35h, 48h, 73h after imbibition).

In order to determine the two different *ANAC060* splice variants two different sets of primers have been used. Primer pair L4 only detects A form, S1 only detects B form to detect both forms we have used primer pair LS3.

Primers details can be found in Table S5. For all analyses the expression was normalized by the expression of two reference genes that are stably expressed in dry seeds: *At4g12590* and *At4g34270* (39).

### Endogenous ABA extraction, detection and external ABA sensitivity analysis

#### ABA extraction and detection

ABA was extracted from 5mg after-ripened dry Col-0 and *anac060-1, anac060-2* seeds, the detailed protocol of ABA isolation, determination and quantification is described in (40).

#### ABA sensitivity analysis

ABA (Duchefa Biochemie) was dissolved in a 10mM MES (Sigma-Aldrich) buffer to make a 25mM ABA master mix (pH 5.8), and diluted to 0.1, 0.2, 0.4, 0.8 µM respectively.

## Acknowledgments

The authors thank Dr Bas Dekkers for critical reflection on the manuscript, professor Teng Sheng offering plasmid containing native promoter of *ANAC060*. This work is financed by the Dutch Research Council (NWO) research domain Applied and Engineering Sciences, project numbers 11314 and 12951. Shuang Song was supported by a fellowship from the China Scholarship Council (CSC).

