## Supporting information for "*DELAY OF GERMINATION 6*, encoding the *ANAC060* transcription factor, inhibits seed dormancy"

Supplementary Information Text

**Fig. S1**. Germination phenotypes of T-DNA knockout lines that are candidates for the gene underlying the QTL.

**Fig. S2**. Germination phenotypes of mutants in the *ANAC060* homologs.

**Fig. S3**. Cellular localisation of the different *DOG6* alleles.

**Fig.S4**. Expression analyses in the *anac060* knock-down mutant.

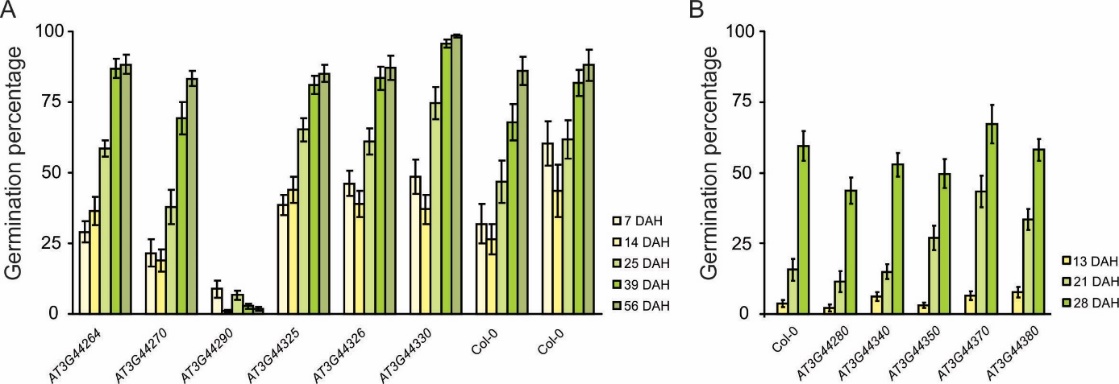

**Fig. S1.** Germination phenotypes of T-DNA knockout lines that are candidates for the gene underlying the QTL. The germination percentage after different days of harvest (DAH) of two independent experiments (A and B) are presented. Details of the KO lines can be found in Table S2.

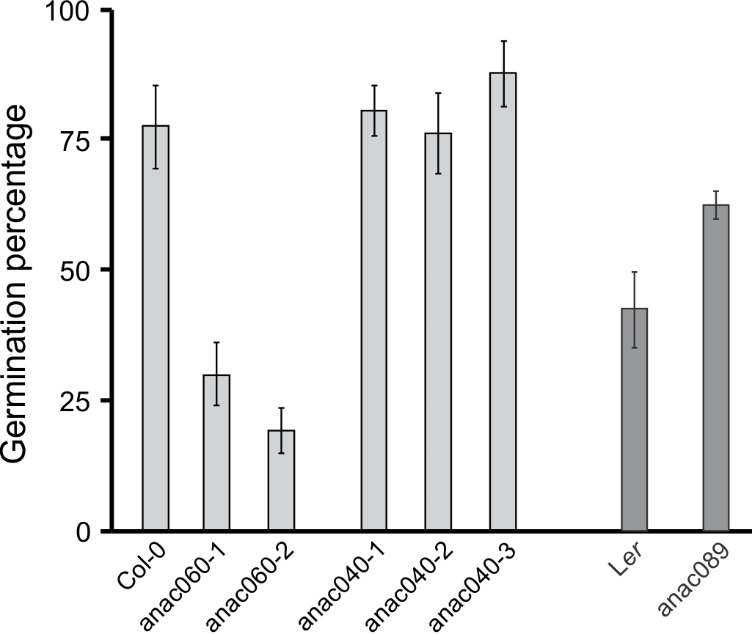

**Figure S2.** Germination phenotypes of mutants in *ANAC060* homologs. The germination percentage of fresh seeds for Col-0 and the different T-DNA lines in the Col-0 genetic background are presented in pale grey, the germination percentage of L*er* and the *ANAC089* KO line (L*er* background) are presented in the darker grey. Details of the KO lines can be found in Table S 2.

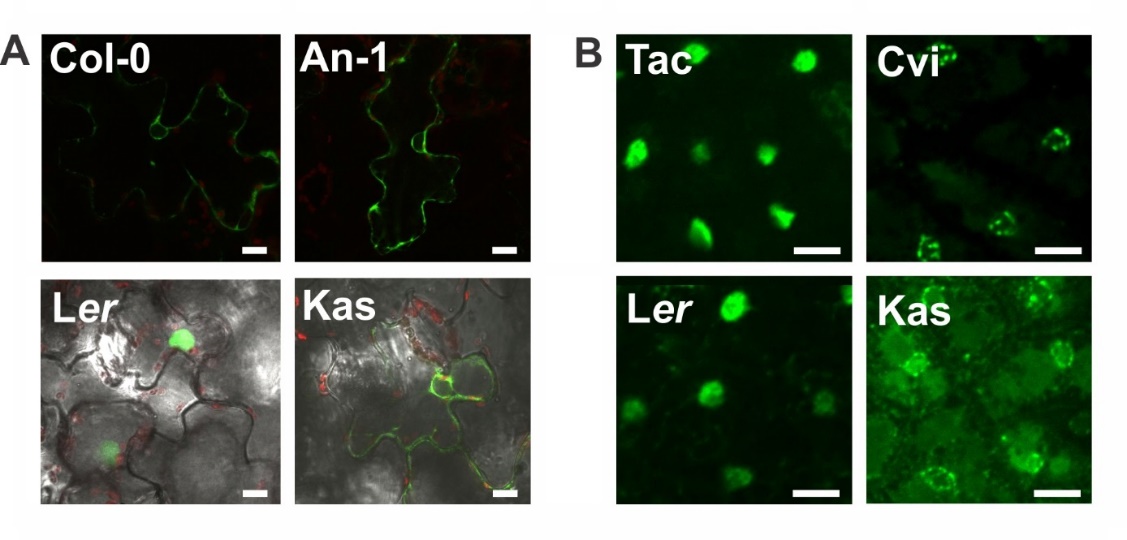

**Figure S3.** Cellular localisation of the different *DOG6* alleles. A. Fluorescence image of GFP fluorescence of the *DOG6*-Col-0, *DOG6*-An-1, *DOG6*-L*er* and the *DOG6* Kas-2 allele in *Nicotiana benthamiana* leaves. B. Fluorescence image of GFP fluorescence of the *DOG6*-Tac, *DOG6*-Cvi, *DOG6*-L*er* and the *DOG6* Kas-2 allele in seeds. The DOG6 protein of the L*er* and Tac allele are localized in the nucleus, for the alleles the localisation is outside the nucleus. Scale bars, 10 µm.

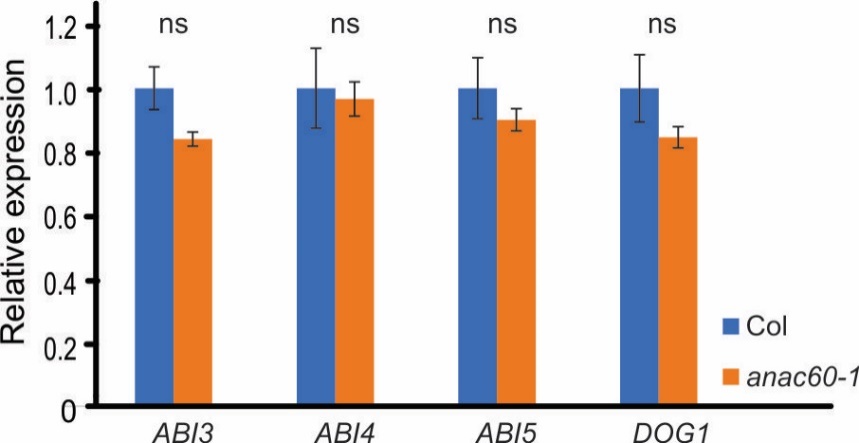

**Figure S4.** Expression analyses in the *anac060* knock-down mutant. *ABI3*, *ABI4*, *ABI5* and *DOG1* expression are not affected in the *anac060-1* knock-down mutant. Relative expression in Col-0 and *anac060-1* is presented. The expression values were normalized using two reference genes (*At2g28390* and *At4g12590*).
